# BiDBiC: A novel ultra-high-throughput pipeline for Bead-in-Droplet Biofilm Cultivation and Characterization

**DOI:** 10.64898/2026.06.18.733193

**Authors:** Jessica D. Li, Xiaoxia N. Lin

**Affiliations:** Department of Microbiology & Immunology, University of Michigan, Ann Arbor, MI; Department of Chemical Engineering, University of Michigan, Ann Arbor, MI

## Abstract

Biofilms are a form of microbial growth consisting of cells, often attached to a surface, embedded in a structured 3D extracellular matrix that confers important emergent properties such as increased resistance to physical removal and antimicrobials. Despite the importance of biofilms to a variety of systems and despite increasing attention from both the public and private sectors, high-throughput approaches to study them are scarce, limiting investigations of complex mechanisms critical for the structure and function of biofilms, such as interactions in multispecies communities. We thus developed a novel workflow to grow and analyze bacterial cells adhered to plastic beads encapsulated within highly parallel nanoliter-scale water-in-oil microfluidic droplets. We term this pipeline for <u>b</u>ead-in-<u>d</u>roplet <u>bi</u>ofilm <u>c</u>ultivation and characterization BiDBiC. To benchmark BiDBiC, we utilized a well-characterized biofilm former, *Stenotrophomonas maltophilia*, as well as a poorly studied drinking water biofilm isolate, *Sphingopyxis* sp. OPL5. Each bacterium exhibited strong adherent growth when co-encapsulated with polystyrene beads in droplets. Furthermore, we retrieved beads from the droplets and removed planktonic cells, enabling focused analysis of adhered cells. From bead-associated biomass, we extracted DNA and RNA for molecular analysis and recovered viable cells for subculturing. We conclude with a discussion of further development of the platform as well as suggestions for microbial biofilm systems that may benefit from ultra-high-throughput droplet-enabled cultivation and analysis.

**Insight Box:** Biofilms are an important yet understudied form of microbial growth. In this study, we developed <u>b</u>ead-in-<u>d</u>roplet <u>bi</u>ofilm <u>c</u>ultivation (BiDBiC), a novel ultra-high-throughput workflow to culture biofilms. The droplets act as massively parallelized miniature bioreactors, with co-encapsulated plastic beads providing a surface for cell attachment and growth. Using a well-characterized biofilm former, *Stenotrophomonas maltophilia*, as well as a drinking water biofilm isolate, *Sphingopyxis* sp. OPL5, we demonstrated robust adherent cell growth in droplets. We additionally efficiently separated beads from planktonic cells, enabling targeted molecular analysis and outgrowth of adherent cells. Adapting and extending BiDBiC could facilitate the study of numerous complex biofilm systems, such as diverse isolates or combinatorial subcommunities of microbiomes, to observe their phenotypes and probe underlying mechanisms.

## Introduction

Biofilms are a ubiquitous and important form of microbial growth (1). They consist of cells embedded within a matrix of extracellular polymeric substances (EPS) composed of polysaccharides, enzymes, and/or DNA, that imparts 3-dimensional structure to the community. This structure gives rise to emergent properties, such as enhanced nutrient utilization and increased antimicrobial resistance, that allow biofilms to colonize and persist in a variety of niches that can have beneficial or detrimental effects on their host or environment (2). For instance, some biofilms in the human body help protect the gastrointestinal tract against invading pathogens while others harbor persistent infections on implanted medical devices; likewise, some biofilms in industrial settings are essential to processes like wastewater treatment while others contribute to biocorrosion of water-exposed surfaces and drinking water quality deterioration (3–7). A better understanding of these biofilms could therefore lead to development of new, effective interventions for controlling and treating pathogenic and undesirable biofilms.

Despite the importance of biofilms, high-throughput approaches for studying them are scarce compared to planktonic cultures. In particular, microfluidic droplets have emerged as an ultra-high-throughput technology platform for the study of various microbial systems, but utilizing them for the investigation of biofilms remains limited due to the inherent complexity of biofilm cultivation and phenotypic characterization (8). One notable previous work, conducted by Niepa *et. al.,* demonstrated generation of droplets with a polydimethylsiloxane (PDMS) shell and successful cultivation of *Pseudomonas aeruginosa* cells in biofilm-like structures within these droplets (9). Surface attachment, a defining feature of many biofilms, however, was not fully enabled by this approach.

We thus developed BiDBiC, an ultra-high-throughput pipeline for <u>b</u>ead-in-<u>d</u>roplet <u>bi</u>ofilm <u>c</u>ultivation and characterization that provides surfaces for cell attachment. In this work, we utilized polystyrene beads as the internal surface to match the plastic typically used in 96-well plates. To benchmark BiDBiC, we utilized an mCherry^+^ strain of *Stenotrophomonas maltophilia*, a well-characterized biofilm former, as well as a robust biofilm-forming isolate from drinking water, *Sphingopyxis* sp. OPL5 (10). Each bacterium exhibited strong adherent growth when co-encapsulated with polystyrene beads in droplets. Furthermore, we retrieved beads from the droplets and removed planktonic cells to enable focused analysis of adhered cells, which remained robustly attached despite chemical and physical treatments. From bead-associated biomass, we extracted DNA and RNA for molecular analysis and recovered viable cells for subculturing. We conclude with a discussion of potential applications of this technology as well as recommendations for further optimization of the pipeline.

## Results

### Overview of the BiDBiC pipeline

We adapted microfluidic water-in-oil droplet technology to develop **BiDBiC**, an ultra-high-throughput pipeline for **<u>b</u>**ead-**i**n-**<u>d</u>**roplet **<u>bi</u>**ofilm **<u>c</u>**ultivation and characterization (Fig. 1). These nanoliter-scale droplets, with a typical diameter in the range of 50-150μm (11), have previously enabled studies of interactions in the vaginal microbiome, high-throughput screening of engineered microbes, and enrichment of rare members of the human gut microbiome (11–13).

**Figure 1.**
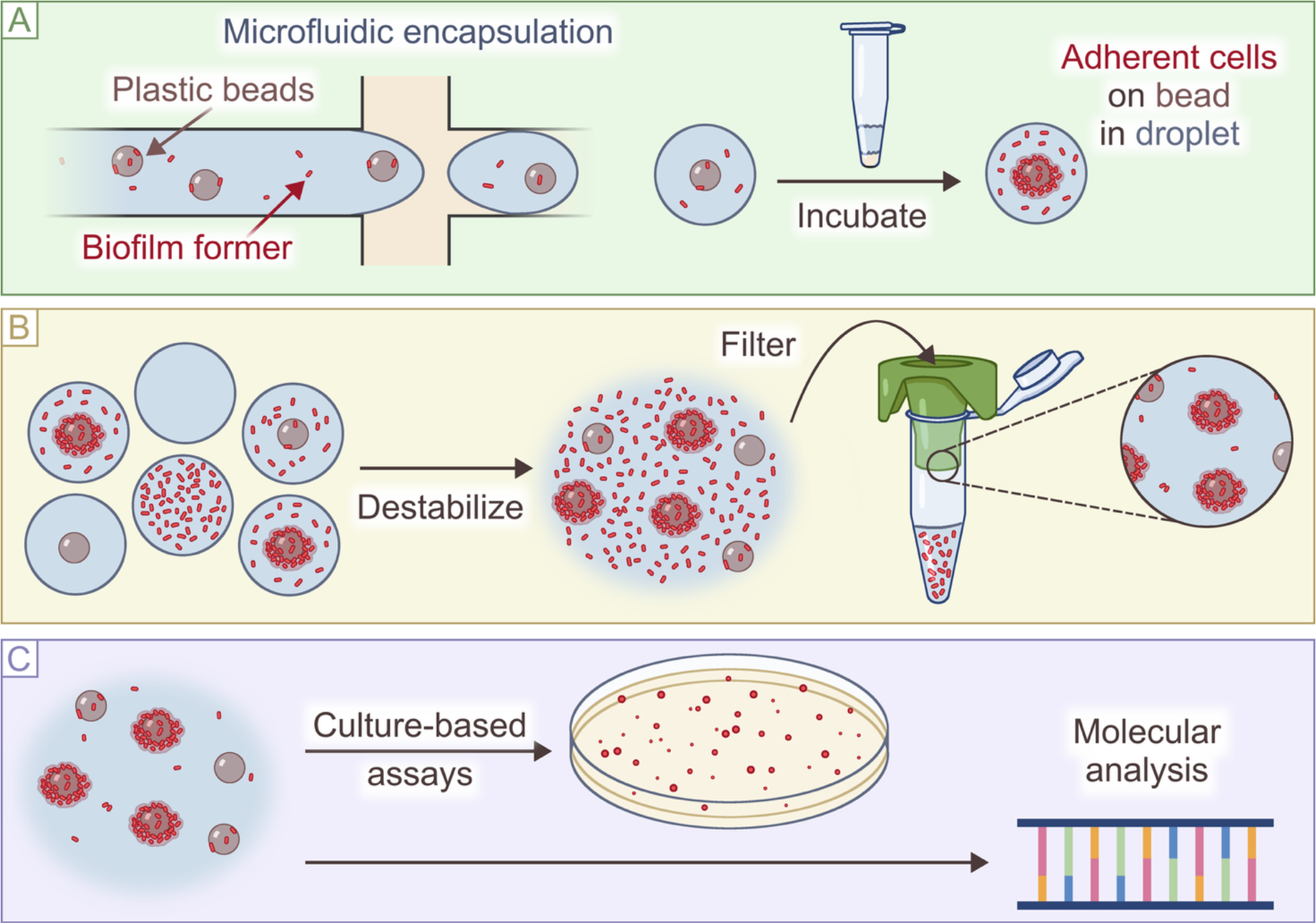
Schematic overview of the microfluidic pipeline for Bead-in-Droplet Biofilm Cultivation and Characterization (BiDBiC). **A**. Plastic beads enable surface-associated growth of biofilm-forming microbes within microfluidic droplets. **B**. Chemical destabilization pools the contents of all the droplets, allowing physical separation of beads (and adhered cells) from planktonic cells for downstream analysis. **C**. Filtered beads can be used for culture-based assays or as template for molecular analysis. Note that the elements of the schematic are not to scale.

Using a microfluidic device with a flow-focusing junction, we generate monodisperse droplets containing mixtures of cells and micron-scale plastic beads. The beads provide solid surfaces for cells to attach to and grow on inside droplets (Fig. 1A). After cultivation, we retrieve cells by destabilizing droplets and aggregating their contents into a single continuous aqueous phase that contain both beads, with adhered cells, and free-swimming planktonic cells. To separate beads from planktonic cells, we filter this aqueous phase (Fig. 1B). The beads with adhered cells can then undergo downstream characterization. In this study, we cultivated cells from beads on agar plates and performed direct molecular analysis of DNA/RNA molecules from bead-adhered cells via PCR and RT-PCR (Fig. 1C).

This platform, in principle, can incorporate beads made of a wide range of materials. Here, we focus on commercially available beads made of polystyrene, which is a plastic used in numerous products including disposable labware.

### Selection of a biofilm-forming bacterium *Stenotrophomonas maltophilia* for validation of BiDBiC

To test and demonstrate the effectiveness of the BiDBiC pipeline, we sought a bacterial species with well-characterized biofilm-forming traits. We selected a strain of *Stenotrophomonas maltophilia* that was previously isolated from drinking water and engineered to constitutively express mCherry for visualization (10). To culture this bacterium, we used Reasoner’s 2A (R2A) medium, a low-nutrient, non-selective medium often used for cultivating slow-growing and stressed bacteria, especially from drinking water samples (14).

We first validated the biofilm-forming property of the *S. maltophilia* strain in R2A on polystyrene surfaces. We started by growing the bacterium in 96-well polystyrene plates without shaking at room temperature (22.5°C), which closely mimics the conditions it experienced in the drinking water system. Under these conditions, wells with *S. maltophilia* retained almost 78 times more crystal violet than the medium-only control after 27 hours of incubation, suggesting the strain forms biofilms robustly (Fig. 2A). We then grew static *S. maltophilia* cultures in 24-well polystyrene plates and imaged washed wells after 1 hour of attachment (0 hours of growth) and after 27 hours of growth. The 1-hour incubation resulted in surface attachment of both single cells and clumps of cells, and after 27 hours we observed significant growth across the entire bottom of the wells with noticeable clumps suggesting 3D structure (Fig. 2B).

**Figure 2.**
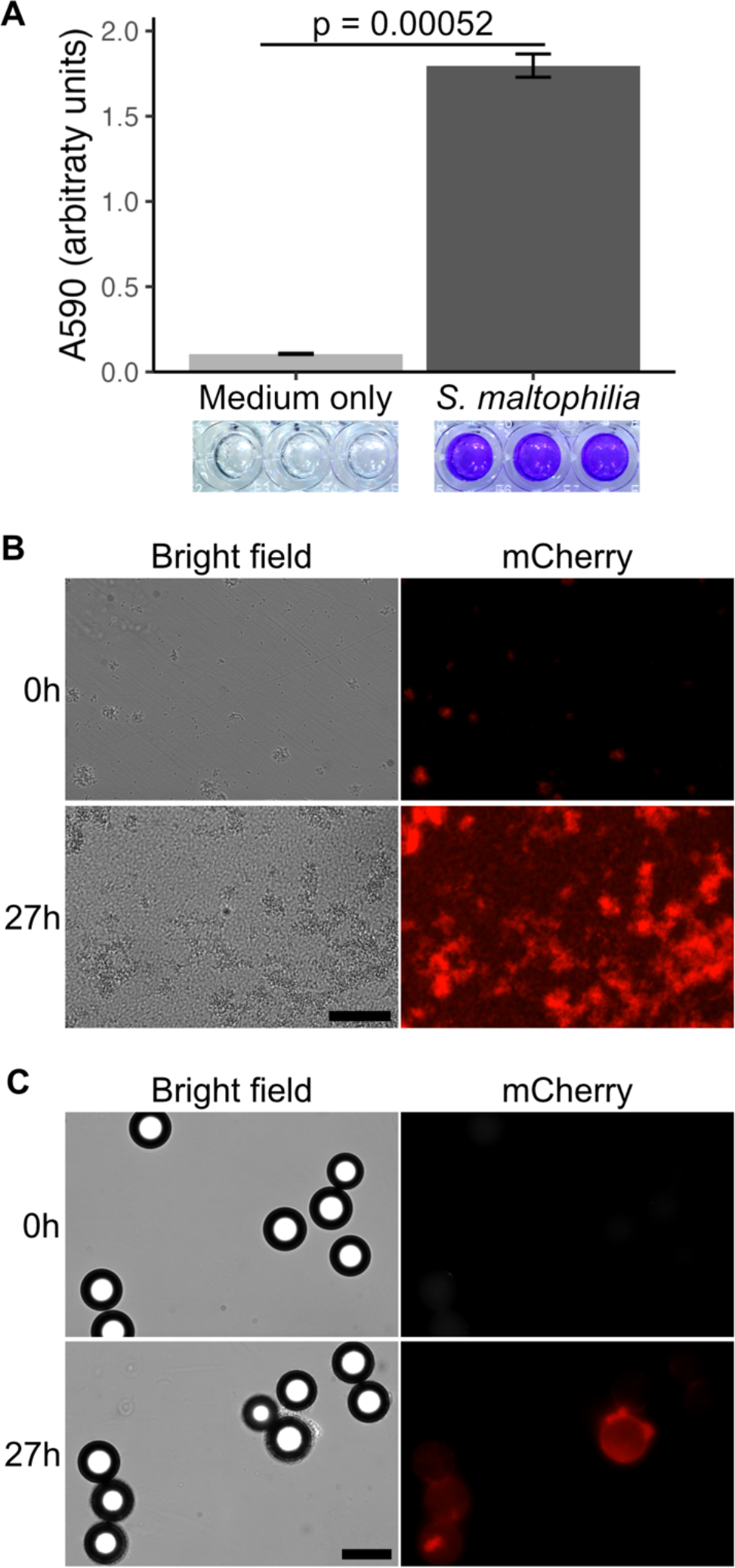
*S. maltophilia* grows robustly on polystyrene surfaces. **A.** Crystal violet staining of 96-well polystyrene plates incubated with medium only (R2A + 20μg/mL gentamicin) or with *S. maltophilia* after 27 hours of static culturing. N = 3 wells/condition, p = 0.00052. **B.** Washed *S. maltophilia* biofilms in 24-well polystyrene plates before and after 27 hours of static culturing. Scale bar = 50μm. **C.** Representative images of 45μm diameter polystyrene beads incubated with *S. maltophilia* for 27 hours. Before imaging, we replaced the medium with sterile PBS. Scale bar = 50μm.

It is important to note that surface geometry can impact biofilm formation and resulting biofilm properties (15,16). Beads that would fit in droplets begin to approach the same length scale as microbial cells, potentially placing constraints on cell attachment due to surface curvature. To determine the impact of bead curvature on *S. maltophilia* cell attachment, we cultured *S. maltophilia* with 15, 25, 45, and 75µm-diameter beads in static milliliter-scale bulk cultures. The beads are denser than water and settle when left undisturbed. To minimize the impact of the oxygen gradient, we incubated the beads in a monolayer in a well plate. When presented with all 4 sizes of beads simultaneously, *S. maltophilia* attached to and grew in biofilm-like, spatially aggregated, dense structures on all sizes of beads (Fig. S1). The channel height of our microfluidic device is ∼80µm, preventing the use of the 75µm beads. We thus proceeded with 45µm beads (Fig. 2C), which provide the largest possible surface for supporting biofilm growth and facilitate separation of bead-adhered cells from planktonic cells for downstream analysis.

### Co-encapsulated beads provide surfaces for adherent *S. maltophilia* growth in droplets

After validating that *S. maltophilia* can grow on 45µm polystyrene beads, we proceeded to test the BiDBiC pipeline. We modified the Drop-seq device developed by Macosko et. al., which was designed to encapsulate resin beads, to utilize a single aqueous inlet flow for simplified operation (Fig. 3A) (17). Because the polystyrene beads are denser than the culture medium, they tend to move slower than the bulk aqueous phase during droplet generation. To more evenly distribute the beads and reduce clogging issues, we loaded the bead-containing aqueous phase into the inlet tubing in small batches separated by pockets of air (Fig. 3B).

**Figure 3.**
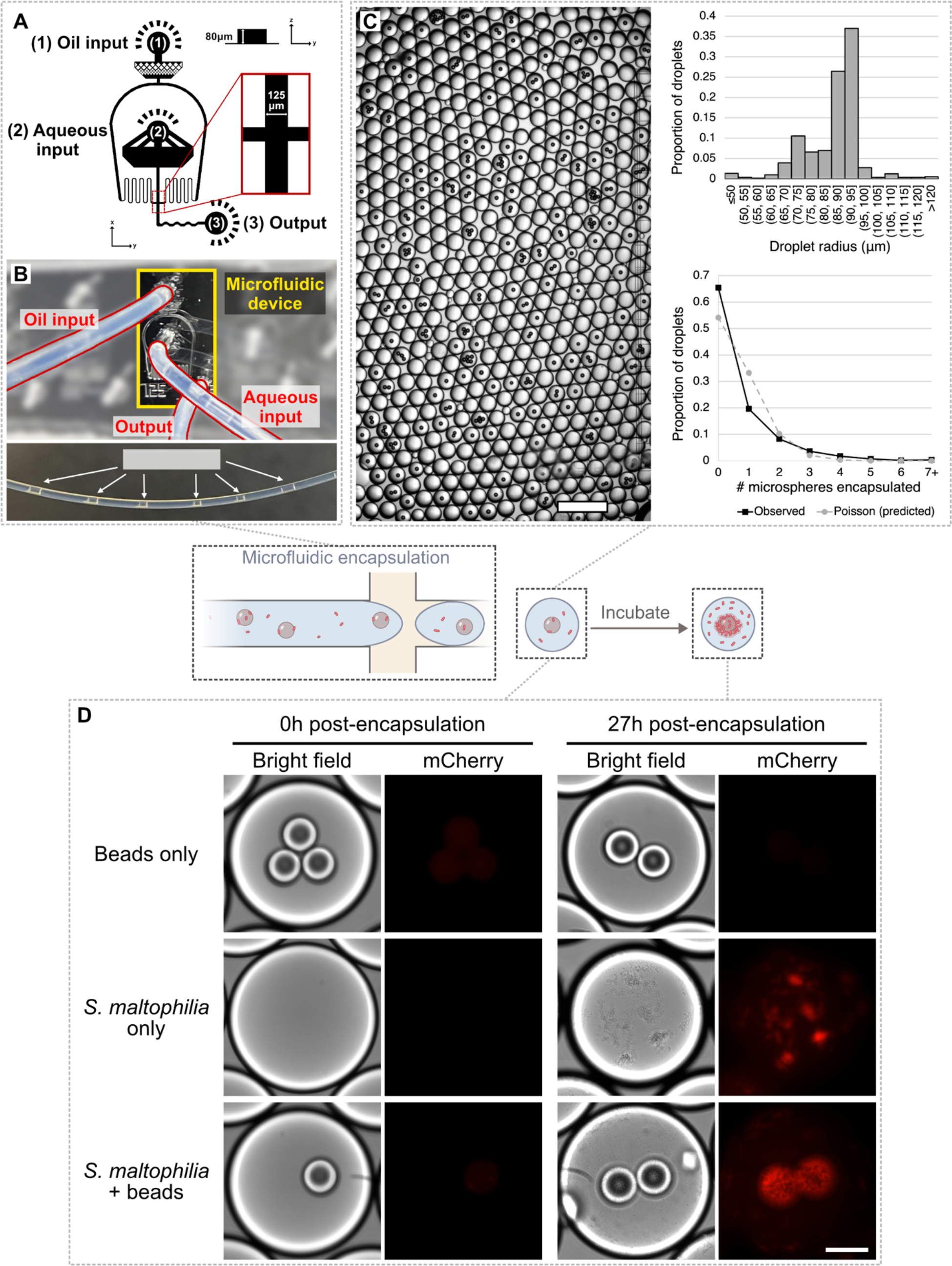
Encapsulated beads provide a surface for cell attachment and adherent growth in microfluidic droplets. **A.** Schematic of the microfluidic device. Portions not fully to scale. Relevant dimensions are labeled. The oil phase enters through inlet (1) and aqueous phase enters through inlet (2). Aqueous droplets pinch off at the flow focusing junction, magnified in the inset, and exit the device through the outlet (3). **B.** Top: picture of droplet generation setup with key components highlighted and labeled. Bottom: picture of tubing for the aqueous input containing the sample in small batches separated by plugs of air, controlled by a syringe (not shown). **C**. Representative image of droplets co-encapsulating beads and *S. maltophilia* cells, distributions of droplet size (N = 1119) and of encapsulated beads per droplet (N = 1651). The mean droplet diameter is 172µm with a standard deviation of 25µm. 34.6% of droplets contain at least 1 bead compared to the 45.9% predicted by the Poisson distribution with λ=0.614 fitted to account for all observed beads. Scale bar = 0.5mm. Equivalent analyses of the bead- and microbe-only droplets are in Fig. S2. **D**. Representative droplets containing 45μm beads only, *S. maltophilia* only, or beads and cells, at 0 and 27 hours after encapsulation and incubation. Images at the two times points do not depict the same droplet but were sampled from the same batch of droplets. Scale bar = 50μm. Additional images are available in Fig. S3.

In one representative experiment, we co-encapsulated beads with *S. maltophilia* cells using this set-up and observed a mean droplet diameter of 172±25µm (Fig. 3C). We also quantified the distribution of beads in droplets. Typically, particle distribution in droplets follows a Poisson distribution with an average of λ particles per droplet determined by the concentration of particles in the aqueous phase (18). Based on the total numbers of observed droplets and beads, we calculated the λ value to be 0.61. The observed distribution deviated slightly from the Poisson distribution with this λ value, however, with fewer droplets than expected containing beads and more than expected containing higher numbers of beads (Fig. 3C). This was likely caused by the tendency of the polystyrene beads to clump together due to increased surface-mediated forces like hydrophobic attraction. Overall, ∼34% of droplets contained at least one bead. We also conducted controls where either beads or cells only were encapsulated and observed similar distributions of droplet size or encapsulated beads per droplet (Fig. S2).

For each condition, we collected ∼400μL of generated droplets in 2mL Eppendorf tubes. We then incubated the droplets at 22.5°C without agitation in the collection tubes. After 27 hours of incubation, we took droplet samples from the Eppendorf tube and examined them via fluorescence microscopy. The beads showed little auto-fluorescence in the red wavelengths, allowing us to track cell growth by visualizing mCherry. As illustrated in Fig. 3D and Fig. S3, *S. maltophilia* displayed robust attachment and growth on beads in the droplets; numerous *S. maltophilia* cells were visible in spatially aggregated, dense structures physically connected to the bead surfaces, in a similar manner as observed in bulk cultures.

### *S. maltophilia* cells remain adhered to beads after droplet pooling and filtration

We noted that droplets contained significant amounts of planktonic cells regardless of whether they contained beads. We thus developed a method to separate beads, with adherent cells, from planktonic cells. First, we chemically destabilized droplets to create a single aqueous phase containing all droplet contents as well as a single oil phase. Following this treatment, a significant portion of beads retained adherent cells in a visually similar attachment pattern as in the droplets and bulk cultures, suggesting the cellular attachments are robust to this chemical and physical treatment (Fig. 4A). We then passed the aqueous phase through a 25µm cell strainer to remove planktonic cells, as this pore size is larger than single cells but smaller than the beads. Again, cells remained adhered to beads in a similar pattern as in the droplets, indicating the physical handling during filtration also does not significantly disrupt cell adhesion (Fig. 4B). Moreover, the density of planktonic cells decreased noticeably after filtration, resulting in significant enrichment of surface-attached cells for downstream analysis (Fig. S4).

**Figure 4.**
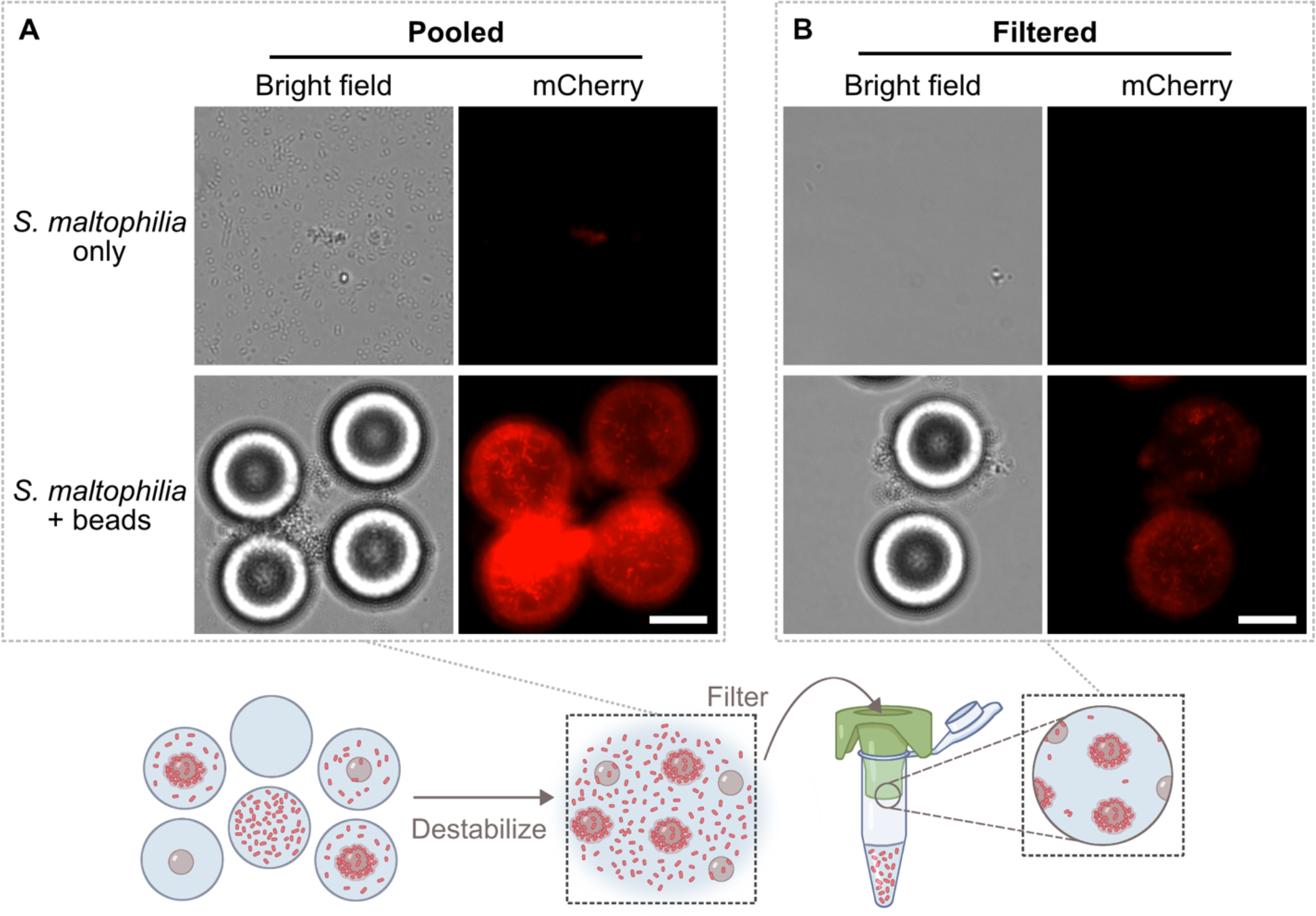
Filtration separates beads from most planktonic cells. **A**. Representative images of *S. maltophilia* cultures after droplet destabilization and before filtration. Scale bar = 25μm. **B**. Representative images of *S. maltophilia* cultures after filtration. Scale bar = 25μm. Additional images are available in Fig. S4.

### Separated beads retain cells suitable for growth-based cellular assays and nucleotide-based molecular analysis

The BiDBiC pipeline currently provides two types of end-point analysis once the beads with adherent cells are retrieved after filtration: growth-based assays that require viable cells to exit the pipeline and molecular analyses that rely on the integrity of DNA and RNA molecules in the cells. We first tested the viability of cells by plating pooled and filtered droplet contents on R2A agar. We utilized a modified dilution series in which we allowed beads to settle for several minutes before plating and diluting samples from the bottom of the tube to ensure a high enough concentration of beads at all dilutions. This did not significantly alter the expected dilution of planktonic cells but allowed us to minimize mechanical disruption of bead-cell adhesion. As shown in Fig. 5A, we observed large colonies centered around beads exhibiting morphologies distinct from those without beads on the same agar or those from *S. maltophilia* cells grown in droplets without beads. Additionally, fluorescence microcopy based on mCherry signal revealed that compared to colonies formed from free cells, the colonies centered around beads displayed highly heterogeneous structures (Fig. 5B). This suggests that the beads carried adherent cells with heterogenous physiology states that seeded the colonies around them.

**Figure 5.**
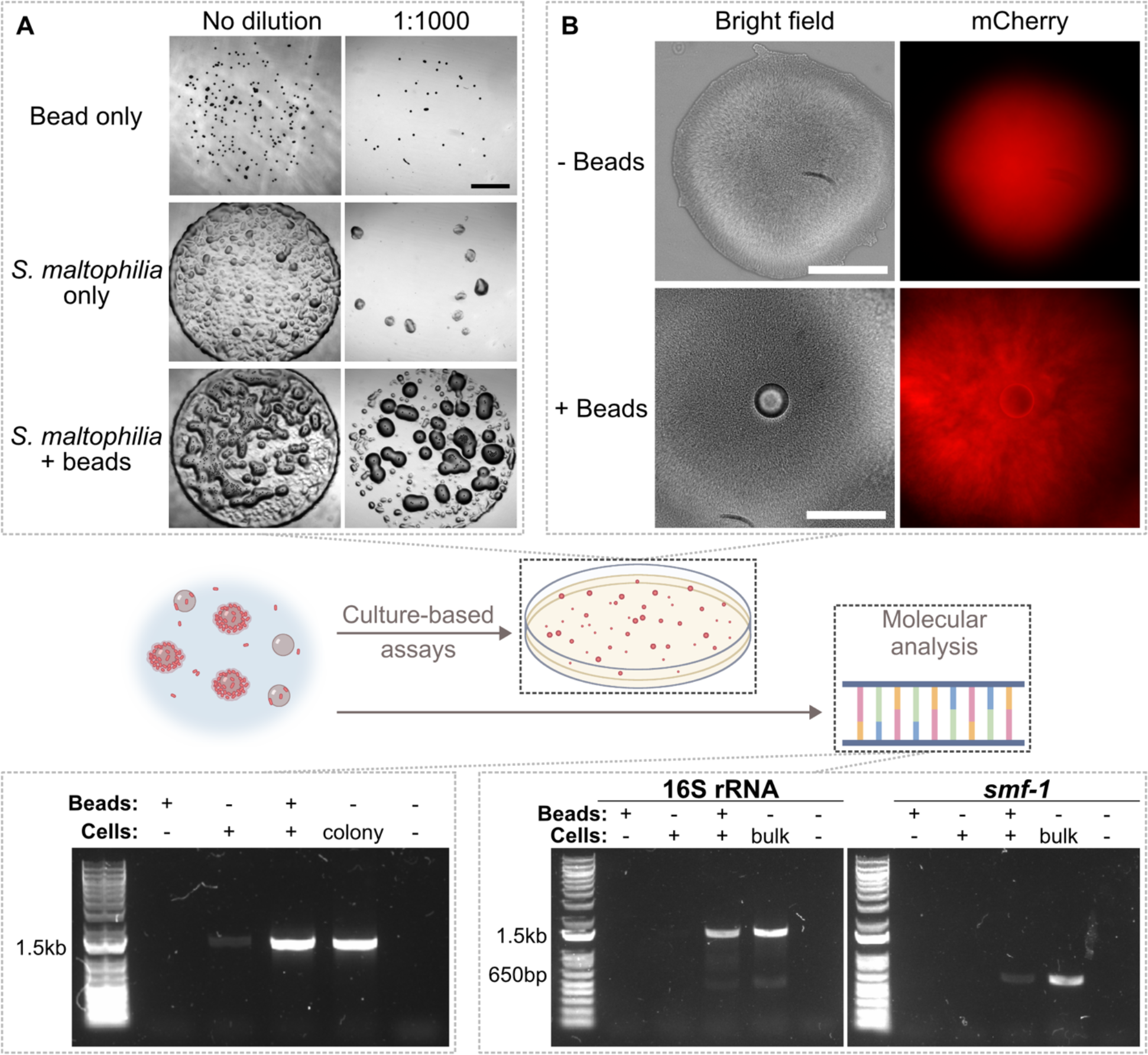
Droplet processing preserves cell viability and nucleic acid quality. **A**. Colony formation on R2A agar from filtered samples after 2 days of incubation at 30°C. Scale bar = 0.5mm. **B**. Close-up of selected colonies from (**A)**. Scale bar = 100μm**. C**. 16S PCR products using filtered samples as template. **D**. 16S rRNA and *smf-1* PCR products using reverse-transcribed cDNA from RNA extractions as template.

Next, we tested the integrity of nucleotides from bead-adherent cells retrieved after the filtration step. This nucleic acid can be subject to molecular assays to provide valuable information such as taxonomic identities via 16S rRNA gene sequencing or active metabolism via RNA analysis. We obtained DNA directly from pooled and filtered droplet contents by lysing sample aliquots with detergent and high heat. Using these samples, we observed strong 16S PCR amplification from the sample that contained both cells and beads, qualitatively only slightly fainter than that obtained from colony PCR (Fig. 5C). In contrast, the PCR reaction utilizing *S. maltophilia* sample that did not contain beads resulted in a barely perceptible DNA band, suggesting most of the amplification in the sample with beads originated from adherent and not planktonic cells. Additionally, we extracted RNA from droplet contents for two-step RT-PCR analysis. We first examined 16S rRNA and again observed clear amplification from cells grown with beads but not without (Fig. 5D).

We then sought to investigate biofilm-associated gene expression and tested four genes previously identified in various *S. maltophilia* strains: *smf-1, rmlA, spgM* and *rpfF* (12). Using primer sets targeting the indicated genes (19,20), we found that the *S. maltophilia* strain used in this study encodes *smf-1* and *spgM*, but not *rpfF* or *rmlA* (Fig. S5). Furthermore, the PCR amplification was particularly strong for *smf-1*, which encodes the fimbriae-1 protein involved in adhesion to both biotic and abiotic surfaces (21) and has been found primarily in the genomes of clinical *S. maltophilia* isolates (20,22–24), despite the fact that the strain used in this study is an environmental isolate. We thus examined *smf-1* expression using RNA extracted from samples retrieved after the filtration step of the BiDBiC pipeline. We were able to detect *smf-1* expression in the sample with beads, though the signal was noticeable fainter than that from high-concentration RNA extracted from a bulk biofilm culture (Fig. 5D). In contrast, there was no detectable amplification from the RNA extracted from droplets without beads, again indicating that the signal detected in the bead-containing sample came from adhered cells. Taken together, our results demonstrate that the expression of an important biofilm-associated gene was maintained in the microenvironment of the droplets and the processing of droplets in the BiDBiC pipeline was able to maintain the integrity of DNA and RNA molecules for downstream analysis.

### Further validation of BiDBiC with a bacterium isolated from drinking water

To assess the general applicability of the BiDBiC pipeline, we tested it with a bacterium we recently isolated from a drinking water biofilm sample. We identified the isolate using 16S rRNA gene sequencing, which had a 100% match with the *Sphingopyxis* sp. OPL5 sequence in the BLAST database (25). This species has only been characterized in a single study, with no prior investigation into its biofilm formation and we verified that the colony and cell morphologies matched either this prior study or closely-related members of the genus (Fig. S6). This “wild” isolate allowed us to assess how compatible the BiDBiC platform is with non-model microbes.

First, we assessed the isolate’s biofilm formation in 96-well culture plates under the same conditions as those used for *S. maltophilia* that mimic indoor plumbing conditions. After 27 hours of incubation, the washed cultures retained almost 45-fold more crystal violet than the medium-only control, indicating significant adherent biomass accumulation (Fig. 6A). We then tested the ability of *Sphingopyxis* sp. OPL5 to adhere to polystyrene beads in bulk cultures. We stained cells with SYBR Green I after 27 hours of incubation to better visualize cell morphologies, and microscopy revealed that cells readily colonized the bead surfaces as spatially aggregated structures, often in a projecting, hairlike arrangement (Fig. 6B, Fig. S7). With this confirmation, we co-encapsulated *Sphingopyxis* sp. OPL5 with 45µm beads in droplets (Fig. 6C). After 27 hours of incubation, we observed distinct attachment of significant amounts of cells to droplets in a pattern similar to their morphology in bulk cultures, in contrast to droplets without beads in which the cells tend to clump together (Fig. 6D, Fig. S8). Thus, beads in droplets likely have the potential to support surface-adhered growth of diverse microbial species.

**Figure 6.**
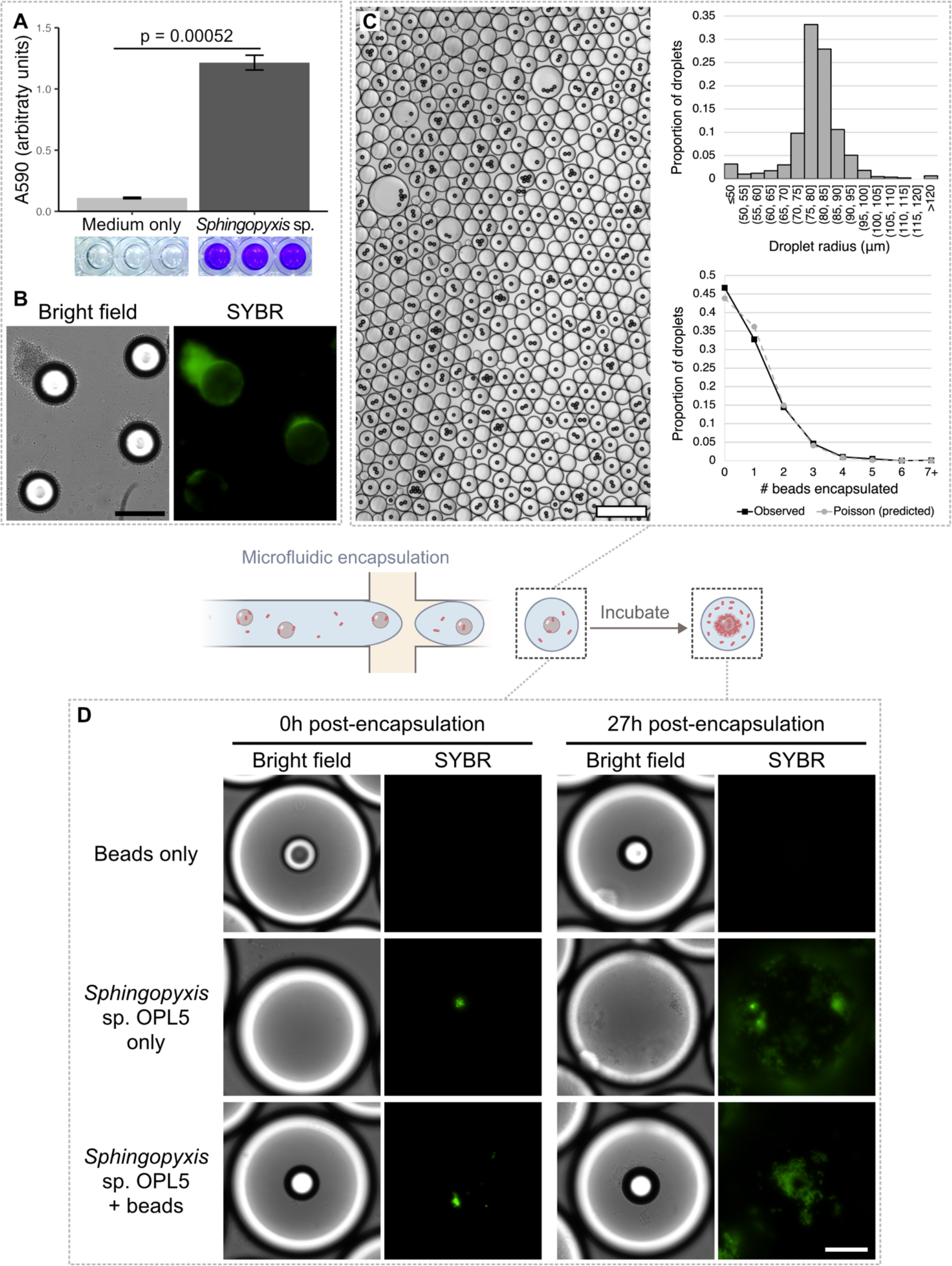
Bead-in-water-in-oil droplets also support growth of a poorly-characterized environmental isolate, *Sphingopyxis* sp. OPL5. **A**. Crystal violet staining of *Sphingopyxis* sp. OPL5 cultures. N=3 for each condition. Standard error for the medium-only datapoints were too low to be visible when graphed. p<0.0005 for *Sphingopyxis* sp. OPL5 compared to medium-only using a 2-tailed Student’s t-test. **B**. Representative microscopy images of *Sphingopyxis* sp. OPL5 bulk cultures incubated with 45μm diameter polystyrene beads for 27 hours. Before imaging at 27h, we replaced the medium with fresh medium. Scale bar = 50μm. **C**. Representative image, distributions of droplet size (N = 1104) and encapsulated beads per droplets (N = 1198) when beads and *Sphingopyxis* sp. OPL5 cells were co-encapsulated. The mean droplet diameter is 158µm with a standard deviation of 24µm. 53.3% of droplets contain at least 1 bead compared to the 56.2% predicted by the Poisson distribution with λ=0.825 fitted to account for all observed beads. Scale bar = 0.5mm. **D**. Representative droplets containing 45μm beads only, *Sphingopyxis* sp. OPL5 only, or beads and cells. Images at the two time points do not depict the same droplet but were sampled from the same batch of droplets. Scale bar = 50μm. Additional images are available in Fig. S8.

## Discussion

In this work, we cultured 3D assemblages of surface-adhered bacterial cells inside water-in-oil microfluidic droplets by co-encapsulating polystyrene beads in an ultra-high-throughput manner. These bead-containing droplets could readily be applied to answer a variety of questions related to surface-associated biofilms in ways similar to prior work from our lab and others that have leveraged droplets to study planktonic microbial systems; for instance, to identify interactions in communities, isolate and study previously uncultured members of microbiomes, and screen bacterial libraries for improved cell activities (11–13). Although we only encapsulated polystyrene beads, the strategy of encapsulating particles within droplets can be extended to other materials for cell attachment provided that the material of interest can be shaped into beads of appropriate sizes for encapsulation in droplets.

Development of BiDBiC is in its early stages. Several improvements are highly desirable and will need future work. As noted earlier, the ∼34% encapsulation efficiency leaves many droplets without a surface for colonization, essentially becoming “wasted” bioreactors. We attribute the deviation of encapsulation from the expected Poisson distribution to two major reasons. First, while we took advantage of bead settling to remove supernatant, some beads are still lost during the washing process after initial cell attachment, leading to a lower concentration of beads than expected and thus a lower λ. If we adjust the λ of the expected distribution so that the total number of expected encapsulated beads per droplets counted matches that of the observed, we see that the number of bead-containing droplets (∼34%) is closer to expected (∼46% vs. ∼53%) (Fig. 3C). Second, the mismatch in density results in non-random distribution of the beads in the aqueous phase. This leads to certain portions of the loaded sample containing low concentrations of beads (lower effective λ, with more empty droplets) and other portions with high concentrations (higher effective λ, with more droplets containing higher numbers of beads).

One avenue to address this issue is to optimize the generation process. We attempted to modify the density of the medium with OptiPrep to more closely resemble that of the polystyrene beads, but this required a large amount of OptiPrep (∼13% v/v) that significantly reduced biofilm biomass measured by crystal violet staining in certain species tested (data not shown). Increasing the concentration (λ) could also reduce the number of empty droplets, but as their concentration increases, the 45µm beads tend to clog the microfluidic device more often. If bead curvature is not a serious issue for a given biological system, decreasing bead size helps as well; we achieved ∼88% encapsulation efficiency at a higher λ in smaller droplets using 25µm beads and device with narrower channels (data not shown). Changes to the device design or using encapsulation methods that do not rely on syringe pumps may also be beneficial. Alternatively, if one has enough cells to generate a larger number of droplets, a droplet sorter may be employed immediately after generation to select only those droplets with beads in them for culturing and analysis, reducing the noise contributed by planktonic cells.

The downstream separation of beads from planktonic cells is another process that would benefit from additional optimization. Since the number of colonies not associated with a bead decreases as expected when diluted, additional rounds of washing may reduce the fraction of planktonic cells in the processed samples. However, this purity would come with the tradeoff of increased cell detachment and sample degradation as the number of liquid handling steps increases. A nontrivial portion of beads also remain trapped in the strainer we used despite several rounds of washing. The use of a custom setup in which the mesh can be detached and fully immersed in buffer to remove more beads may improve the sample yield.

The last major limitation is the extent to which the bead-in-water-in-oil system can represent the conditions biofilms undergo. This issue is not unique to our methodology; all models have their limitations in terms of physiological or biological relevance. Thus, before applying our methodology to study a given system, we believe there are two major factors to consider. First is the issue of surface curvature. The species we tested in this work appear to attach well to a variety of sizes of beads with a range of surface curvatures, but this may not be true for all species. One way to overcome this issue is to use flattened particles instead of spherical ones, though this requires additional labor and equipment as well as potential modification of the encapsulation protocol. The second issue is that there is no fluid flow in the droplets. This can affect biofilm development, which often relies on physical cues such as shear stress, and limits the droplets to represent batch cultures, in which cell growth is limited by depletion of nutrients and accumulation of waste.

We thus suggest utilizing our bead-in-droplet system as an initial screen of large search spaces of biofilm-forming microbial strains or communities, followed by complementary methodologies at larger scales to more closely examine interesting candidates. The ultra-high-throughput nature of droplet generation and encapsulation is ideal to assess, for instance, the phenotypes of single cells from libraries or to rapidly generate random combinations of members from complex mixed-species samples. Combined with fluorescence reporting through genetic modification or probes, we could sort for droplets containing cells with desirable growth or activity (11,26). We could then grow up cells retrieved from these sorted droplets for additional assays at microfluidic or conventional milliliter lab scale that are lower throughput but would more accurately represent the conditions biofilms face (27). Additionally, we and others have utilized microfluidic droplets to enrich for previously uncultured microbial species, another application that could be extended to biofilm and surface-associated communities (13).

Certain biofilm communities, particularly those centered naturally around small particles or those in stagnant conditions, may be particularly suitable for study using our methodology. Since we utilized polystyrene beads for our proof-of-concept work, our methodology could help accelerate studies on the effect of community membership and interspecific interactions in microbe-mediated degradation of microplastics. From soil samples, cultivation of single cells with beads may allow us to isolate new species, and co-cultivation with fluorescent pathogens of interest may aid in ultra-high-throughput novel antimicrobial discovery. Our success with two drinking-water-derived bacteria in this work also suggests our bead cultivation strategy could represent areas of stagnant water or high retention time in the drinking water distribution system, allowing us to study how individual isolates or consortia derived from drinking water biofilms respond to such conditions.

As microfluidic experimental techniques gain traction in the field of microbiology, our work seeks to expand the applications of microfluidics into new sub-disciplines and provide researchers with a more labor- and resource-efficient method of performing ultra-high-throughput biofilm cultivation. With the limitations and suggestions laid out above, we hope to provide the necessary information for researchers to make informed decisions on the suitability of our methodology to their systems and questions of interest.

## Materials and Methods

### Lithography

We generated SU-8 master molds of the microfluidic device (Fig. 3A) using a standard SU-8 photolithography protocol in a cleanroom. A 4-inch diameter silicon wafer was spincoated with SU-8 2050 (Kayaku Advanced Materials) to an 80µm thickness and baked at 65°C and 95°C for 6 and 18 minutes, respectively. A transparency mask with the design was purchased from Fine Line Imaging, aligned to the wafer, and exposed to UV for 10 seconds following vacuum-assisted contact. Post-UV exposure, the water was baked again at 65°C and 95°C for 4 and 12 minutes, respectively, and developed with SU-8 developer. The wafer was washed multiple times with isopropanol and SU-8 developer before being blow-dried. After profilometer assessment of device thickness, the wafer was hard baked at 175°C for 5 minutes and silanized with 2-3 droplets of tridecafluoro-1,1,2,2-tetrahydrooctyl-1-trichlorosilane by vapor deposition in a vacuum desiccator for 1 hour. Finally, the wafer was baked at 150°C for 10 minutes.

We then performed standard polydimethylsiloxane (PDMS) soft lithography to generate the microfluidic devices. 9 parts PDMS elastomer base was mixed with 1 part curing agent and poured onto the SU-8 master mold taped to the bottom of a large petri dish. The PDMS mixture was degassed in a vacuum chamber and cured for 18 hours at 65°C at atmospheric pressure. Cured PDMS devices were cut with a razor, peeled off the master mold, and cleaned with tape. Inlet and outlet holes were punched out using a 1.0mm biopsy punch, and the PDMS was bonded onto glass microscopy slides with a plasma wand. Devices were then baked at 80°C for 10 minutes and silanized by vapor deposition in a vacuum chamber overnight. Finished devices were covered with tape until use.

### Microbial cultures

We utilized a *S. maltophilia* strain transformed with plasmid pBPF-mCherry encoding gentamicin resistance (10) and a *Sphingopyxis* sp. OPL5 strain isolated in this study (Fig. S6). R2A (Neogen NCM0188A) containing 0.5g/L of yeast extract, meat peptone, casamino acid, glucose, and starch; 0.3g/L of K_2_HPO_4_ and sodium pyruvate; and 0.05g/L of MgSO_4_ was used as the culture medium. All *S. maltophilia* liquid cultures and agar plates contained 20µg/mL gentamicin. For *S. maltophilia*, 2mL R2A precultures were inoculated with a single colony picked from 1.5% agar R2A plates streaked with 25% glycerol cryostocks. For *Sphingopyxis* sp. OPL5, 2mL R2A precultures were directly inoculated from cryostocks. Agar plates were parafilmed and incubated at 30°C while precultures were incubated at 30°C overnight (∼12-16 hours) with shaking at 250rpm. Cells from precultures were washed 3 times with R2A by centrifugation at 4000xG for 5 minutes, resuspended in fresh medium, and diluted as necessary for counting in a Neubauer Improved (NI) C-Chip hemocytometer (INCYTO DHCN015) before use in assays.

### Crystal violet assay

We performed a crystal violet stain as previously described, with slight modifications, to assay biofilm biomass (28). Briefly, we diluted overnight precultures to an OD600 of 0.005 and plated 100µL/well of sterile medium or cells in 96-well non-treated polystyrene plates (VWR 10861-562). We incubated plates at 22.5° C without shaking for 27 hours before readout. To remove planktonic cells, we removed the supernatant and washed once with 200µL/well sterile PBS. We then stained the remaining biomass by incubating wells with 200µL 0.2% (w/v) crystal violet at room temperature for 15 minutes. After removing the stain, we washed the wells 4 times with 200µL PBS and allowed the plates to air dry for 20 minutes at room temperature with the lid removed. To elute retained crystal violet, we incubated wells with 95% (v/v) ethanol for 20 minutes at room temperature before transferring 150µL to a new plate and reading out the absorbance at 590nm in a BioTek Synergy H1 plate reader.

### Preparation of beads

We purchased 15, 25, 45, and 75µm diameter uncoated polystyrene beads crosslinked with divinylbenzene from Polysciences Inc. (catalog numbers 18328-5, 07313-5, 07314-5, and 24049-5 respectively). Bottles were shaken to evenly disperse beads before 200-500µL was squeezed into 1.5µL Eppendorf tubes. Beads were sterilized by washing three times with an equivalent volume of 70% ethanol at 3000xG centrifugation for 3 minutes and were then washed 4 times with an equivalent volume of R2A before suspension in a final volume of 100-200µL R2A. When necessary, washed beads were diluted by up to 10 times in R2A before counting in a NI C-Chip hemocytometer.

### SYBR Green I staining

10,000x concentrated SYBR Green I (Invitrogen S7563) was diluted 1,000x in Novec HFE-7500 Engineered Fluid and stored at -80°C. 2μL stain was added to 8μL of droplets in a PCR tube, incubated for 5 minutes in the dark at room temperature, and gently inverted before loading in an NI C-Chip hemocytometer.

### Biofilm morphology assay

24-well non-treated polystyrene plates (CELLTREAT 229524) were inoculated with 500µL of 10^7^/mL prepared cells and incubated statically for 1 hour at room temperature to allow cells to attach. Wells were then washed once with R2A and replenished with 500µL of fresh R2A before 27-hour incubation at 22.5°C. After the second incubation, wells were washed once with sterile PBS and replenished with 500µL PBS for imaging. *Sphingopyxis* sp. OPL5 wells were additionally stained with SYBR Green I prior to imaging to visualize cells.

### Cell attachment to beads

Washed cells and beads were either combined at 10^7^/mL (λ=23.5) and 3.18x10^5^/mL (λ=0.75) respectively or diluted to those concentrations separately. Prepared cells and beads were incubated at room temperature for 1 hour to allow cells to attach to the surface before 90% of the supernatant was carefully removed and fresh medium added. This wash was repeated once after giving the beads enough time to settle back to the bottom. Control samples without beads were handled in an identical manner as those with beads.

### Bulk culturing of cells with beads

500µL of prepared cells and/or beads were added to 24-well non-treated polystyrene plates and incubated without shaking at 22.5°C for 27 hours. After incubation, the supernatant was gently aspirate and pipetted over the bottom of the well at an angle to dislodge beads before transfer to a 1.5mL tube. The tube was pulse centrifuged at low speed and most of the supernatant was removed. The well was washed twice with 500µL PBS, the PBS was transferred to the tube each time, and most of the supernatant was removed after pulse centrifugation before 500µL sterile PBS was added to the washed beads. *Sphingopyxis* sp. OPL5 samples were stained with SYBR Green I. Settled beads were collected from the bottom of the tube for imaging in a NI C-Chip hemocytometer.

### Co-encapsulation of cells and beads

Kent Scientific GenieTouch syringe pumps controlled the flow of the oil (30µL/min) and aqueous (20µL/min) phases during droplet generation. The oil phase consisted of Novec HFE-7500 with 2% w/v PEG-PFPE amphiphilic block copolymer surfactant (Ran Biotechnologies, 008-FluoroSurfactant) loaded in a 3mL syringe with a 24-gauge Luer Lock syringe needle connected to a length of PTFE tubing (0.022” ID, Cole-Parmer). To load the aqueous phases of prepared cells and/or beads, 1mL syringes were first filled with R2A and connected to 24-gauge needles and PTFE tubing. R2A was manually pushed through the entire length of tubing before the syringe was secured in the syringe pump. Aqueous phases were drawn up in 2.5-5µL increments separated by pockets of air using the manual withdrawal button.

Microdroplets were collected directly from the outlet tubing into 2mL Eppendorf tubes and incubated at 22.5°C without agitation.

### Droplet pooling

360µL droplets were combined with an equal volume of Novec HFE-7500 with 20% 1H,1H,2H,2H-Perfluoro-1-octanol and mixed by gentle flicking. The tubes were pulse centrifuged at low speed to separate the oil and aqueous phases. As much of the denser oil phase as possible was removed before an equal volume of 1% SPAN-80 in hexane was added and mixed by gentle flicking, followed by pulse centrifugation at low speed. As much hexane as possible was removed and an equal volume of sterile PBS was added.

### Bead filtration

20µm pluriStrainer cell strainers (Pluriselect 43-10020-50) were washed with 100µL sterile PBS and placed in 2mL tubes. The pooled droplet contents were added to the strainer and gently washed 5 times with 360µL PBS, and excess liquid was pipetted off the bottom of the strainer after each wash. The strainer was then turned upside down over a fresh 1.5mL tube, and 3 washes of 150µL of PBS was pipetted onto the bottom of the strainer and allowed to flow through. Excess liquid was carefully pipetted from the inside of the stainer each time. 400µL supernatant was removed from the top after pulse centrifugation at low speed and replaced with fresh PBS twice to further dilute the planktonic fraction.

### Droplet content plating

The pooled and filtered droplet contents were given enough time for the beads to allowed to settle before taking aliquots from the bottom for serial 1:10 dilutions in R2A. Samples without beads were diluted in an identical manner to those with. 5µL were spot plated on R2A with 1.5% agar in individual wells of 24-well plates and allowed to air dry before incubation at 30°C for 2 days.

### Microscopy and image processing

Phase contrast and fluorescence microscopy were performed with a Nikon Eclipse Ti-S inverted light microscope equipped with a CoolLED pE-300lite white light LED illumination system, and images were captured with Ocular advanced scientific camera control software version 2.0 using a QImaging Retiga R6 Monochrome CCD camera, except for Gram staining pictures, which were captured with an iPhone SE through the eyepiece. mCherry images were taken with the TRITC filter and SYBR Green I images were taken with the EYFP filter. All images were captured with 500ms exposure time and an objective lens of 20x, except for droplet quantification and droplet plating images, which were captured with 15ms exposure time and an objective lens of 2x.

Images were processed in ImageJ (version 1.54g) to increase the brightness of the raw images and decrease background noise, with all images from a given day in the same channel adjusted identically. Color was added to the mCherry and SYBR Green I images by using the overlay blending function in Firealpaca (version 2.13.7) on the raw images.

### PCR amplification

PCR was performed in 12.5µL reactions using a premixed *Taq* PCR master mix (Thermo Scientific K0171) with 1µL of template and 0.2µM of each primer. Table S1 lists the primer sequences and expected fragment sizes. When samples with whole cells were used, they were lysed in a solution containing 10mM Tris-HCl, 0.1% Triton-X, and 1mM EDTA. The initial denaturation step was 5 minutes at 95°C. The amplification cycles consisted of 20 seconds at 95°C, 30 seconds at 50°C, and 2 minutes at 68°C. 25 cycles were used for all amplification except Fig. S1, which was amplified for 30 cycles. Reactions were given 5 minutes at 68°C for final extension and refrigerated at 4°C for up to a day before gel electrophoresis. 4µL of each reaction was run for 25 minutes on a 1% agarose gel at 120V, stained with ethidium bromide, and visualized with UV light. Photos were all taken with a 0.5 second exposure time.

### RNA extraction and reverse transcription

Samples were stored for no longer than 1 month at room temperature in Zymo DNA/RNA Shield. RNA was extracted using the parallel purification protocol of the ZymoBIOMICS DNA/RNA Miniprep Kit with 15 minutes of bead beating and in-column DNase I digestion. All spins were performed at 16,000xG for 30 seconds unless otherwise required by the kit protocol. Extracted RNA was eluted with 50µL buffer. Reverse transcription was performed with the NEB ProtoScript II First Strand cDNA Synthesis Kit using the provided random primer mix. The standard protocol was used with the recommended 5-minute 25°C incubation step between sample denaturation and reverse transcription.

## Acknowledgements

The authors thank Drs. Ameet J. Pinto and Lutgarde Raskin for instrumental feedback throughout the course of this work, Dr. James Y. Tan for invaluable advice, Dr. Chuanwu Xi for providing the *S. maltophilia* strain, and Zymo Research for providing a complimentary Microbiomics - DNA/RNA Graduate Student Starter Pack.

## Data Availability Statement

The data supporting this article are available in the main body and online supplementary material. Additional images are available upon request to the corresponding author.

