## Supplementary materials for "BiDBiC: A novel ultra-high-throughput pipeline for Bead-in-Droplet Biofilm Cultivation and Characterization"

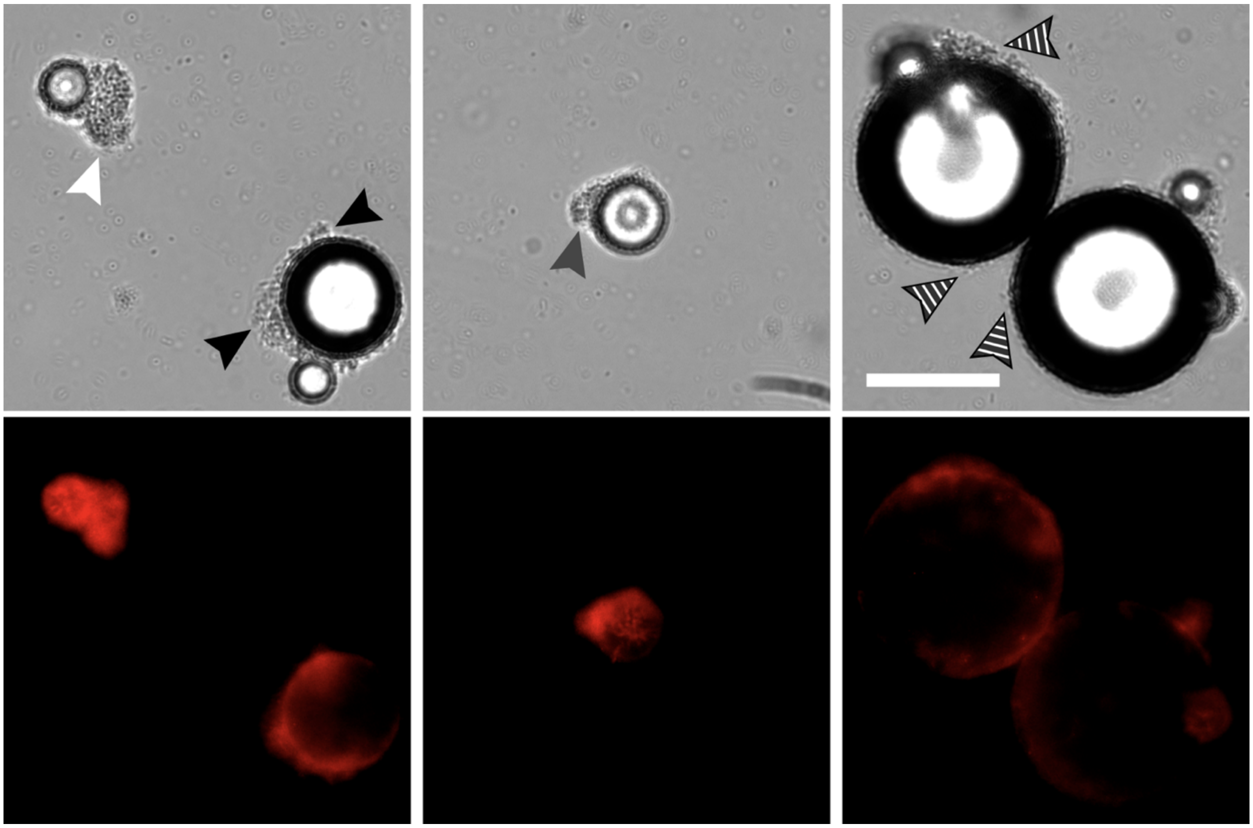

**Figure S1.** ***S. maltophilia*** **cells attached to and grew on 15μm (white arrows), 25μm (gray arrows), 45μm (black arrows), and 75μm (striped arrows) diameter beads after 27 hours of incubation at 22.5˚C.** Scale bars = 50μm.

**Alt text**: Microscopy images of plastic beads and fluorescent cells.

**A**

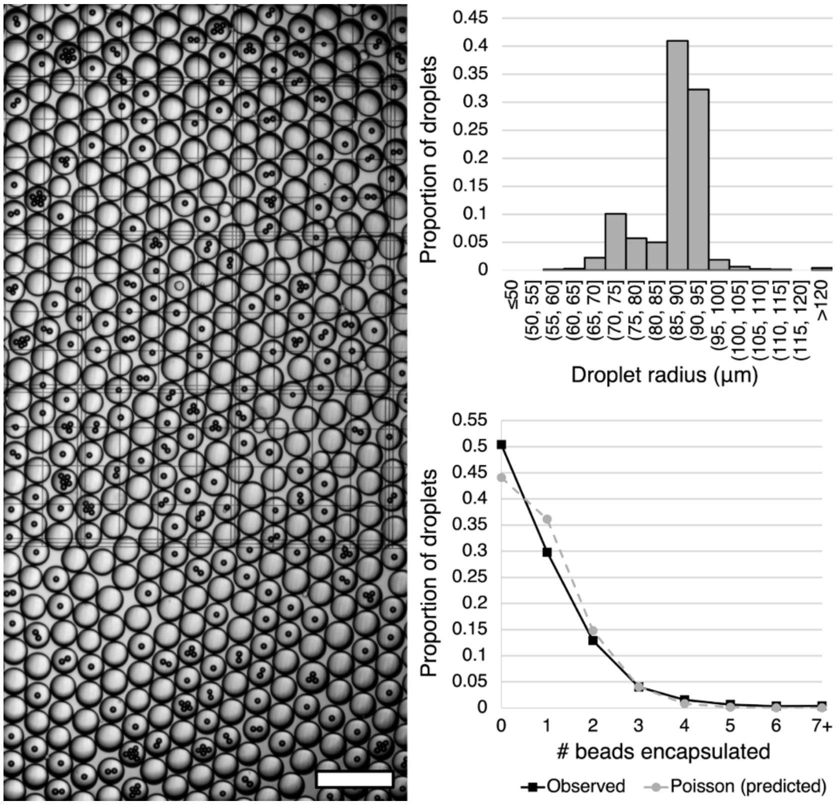
0.45
0.4
0.35
0.3
0.25
0.2
0.15
0.1
0.05
0
Proportion of droplets
≤50
(50, 55]
(55, 60]
(60, 65]
(65, 70]
(70, 75]
(75, 80]
(80, 85]
(85, 90]
(90, 95]
(95, 100]
(100, 105]
(105, 110]
(110, 115]
(115, 120]
>120
Droplet radius (μm)
0.55
0.5
0.45
0.4
0.35
0.3
0.25
0.2
0.15
0.1
0.05
0
Proportion of droplets
0 1 2 3 4 5 6 7+
### beads encapsulated
Observed
Poisson (predicted)

**B**

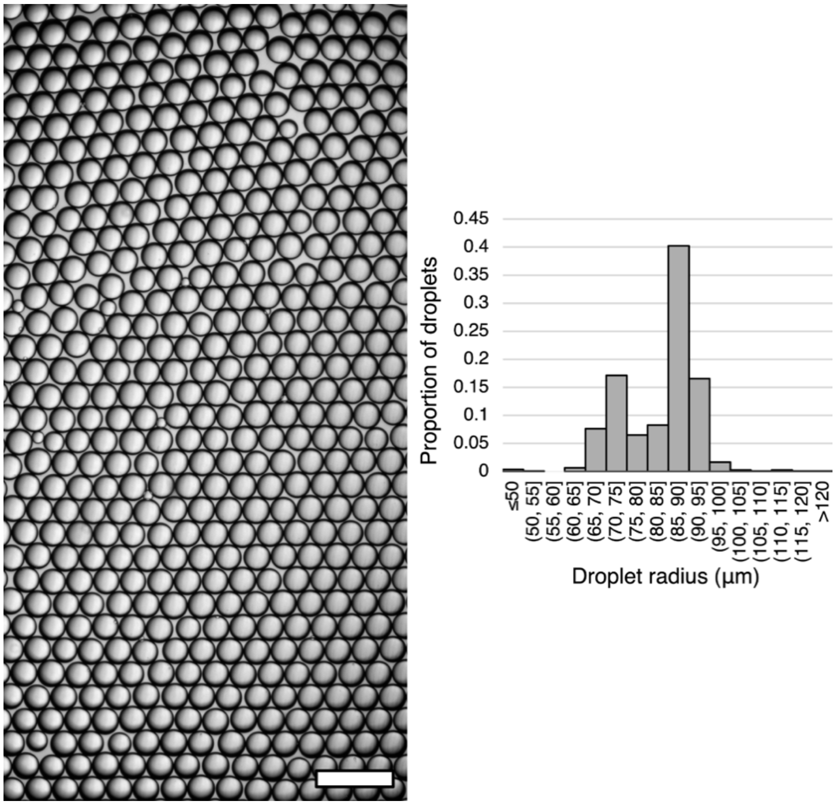
0.45
0.4
0.35
0.3
0.25
0.2
0.15
0.1
0.05
0
Proportion of droplets
≤50
(50, 55]
(55, 60]
(60, 65]
(65, 70]
(70, 75]
(75, 80]
(80, 85]
(85, 90]
(90, 95]
(95, 100]
(100, 105]
(105, 110]
(110, 115]
(115, 120]
>120
Droplet radius (μm)

**Figure S2. Quantification of bead-only and *S. maltophilia*-only droplets. A**. Representative image of droplets containing only beads, distribution of droplet size (N = 2009), and distribution of encapsulated beads per droplet (N = 1382). The mean droplet diameter is 174μm with a standard deviation of 21μm. 49.6% of droplets contain at least 1 bead compared to the 55.9% predicted by the Poisson distribution with $\lambda$=0.819 fitted to account for all observed beads. Scale bar = 0.5mm. **B**. Representative image of droplets containing only *S. maltophilia* cells and distribution of droplet size (N = 1074). The mean droplet diameter is 166±19μm. Scale bar = 0.5mm.

**Alt text**: Microscopy images and graphs quantifying droplet size and bead distribution in droplets.

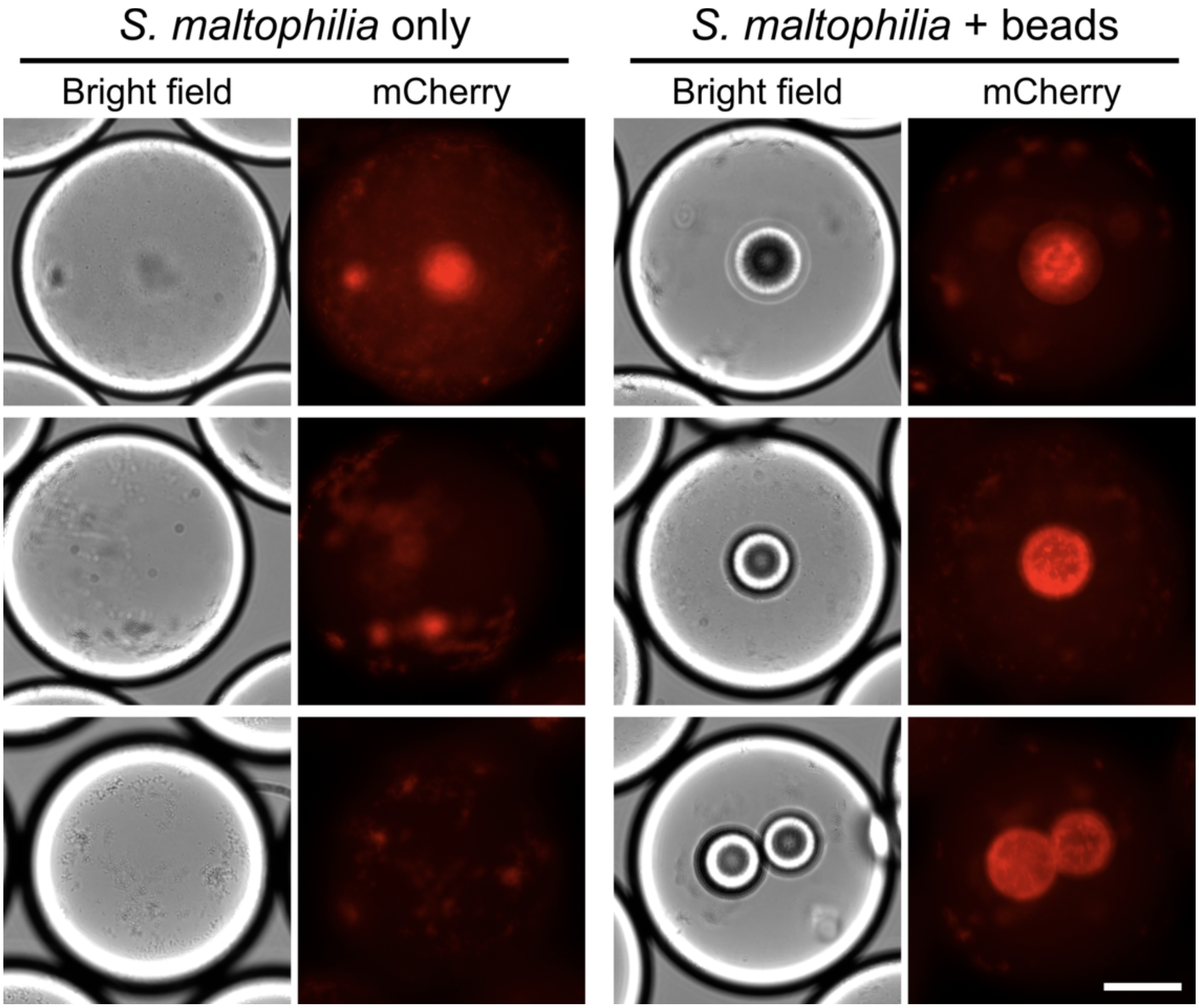


**Figure S3. Additional images of droplets containing *S. maltophilia* with or without beads after 27-hour incubation.** Scale bar = 50μm.

**Alt text**: Zoomed-in microscopy images displaying cell attachment to and growth on the beads in droplets.

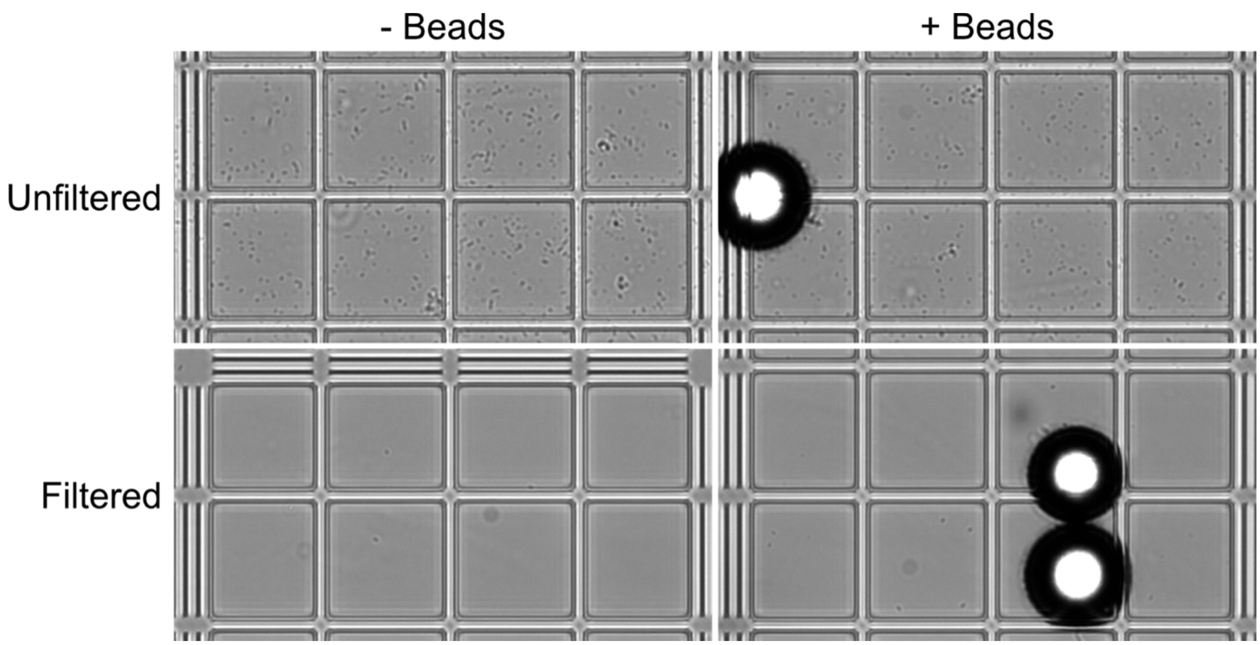


**Figure S4. Planktonic cell densities decreased significantly after filtration, with or without beads.** Each square in the grid is 50μm x 50μm.

**Alt text**: Microscopy images against a gridded background to compare free-swimming cell density across samples and workflow steps.

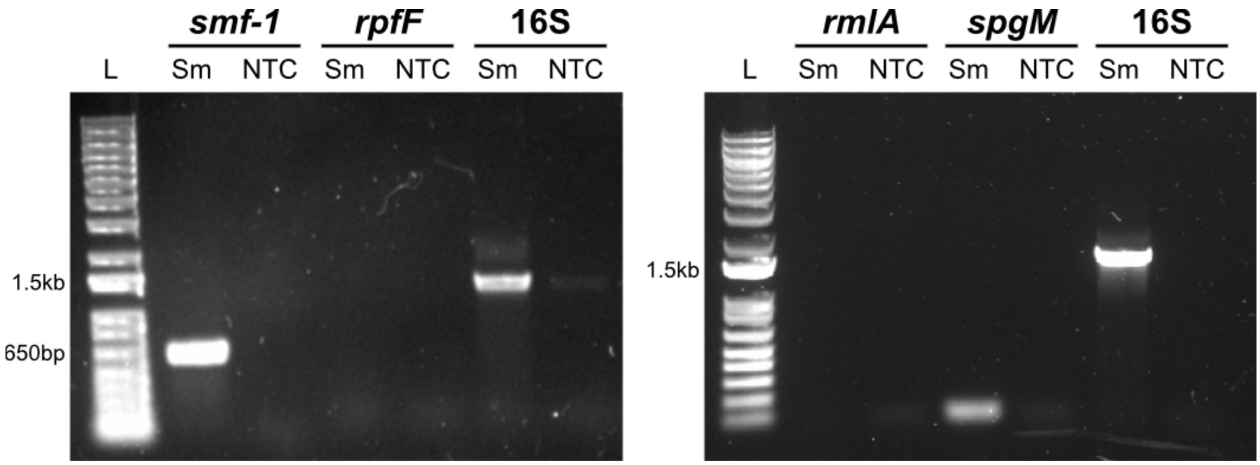


**Figure S5. The *S. maltophilia* strain used in this study encodes biofilm-associated genes *smf-1* and *spgM* but not *rpfF* or *rmlA*.** PCR products using primer sets targeting the indicated biofilm-associated genes (1,2). L = ladder, Sm = *S. maltophilia*, NTC = no template control.

**Alt text**: Gel electrophoresis scans showing which genes the *S. maltophilia* strain in this study expresses.

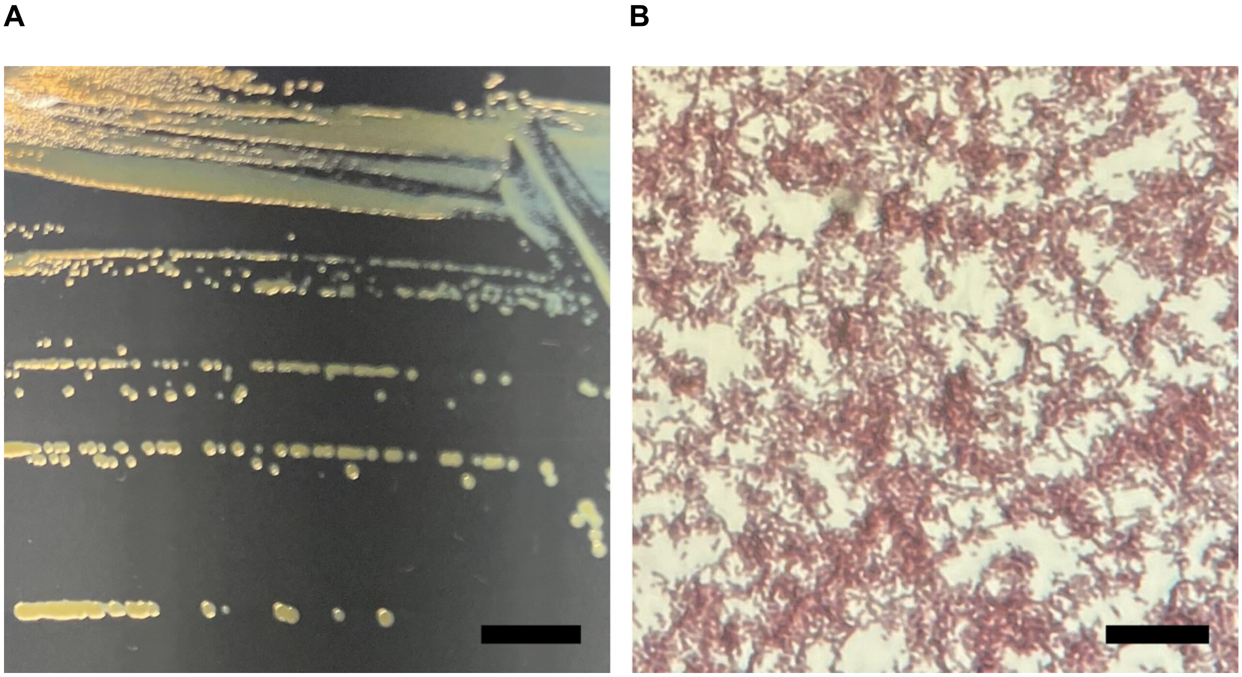

**Figure S6. A drinking water biofilm isolate identified as *Sphingopyxis* sp. OPL5**. **A**. Colony morphology of the isolate. Scale bar = 0.5mm. **B**. Gram staining of cells. Scale bar = 25µm.

**Alt text**: Photograph of colonies on agar and microscopy image of Gram-stained cells.

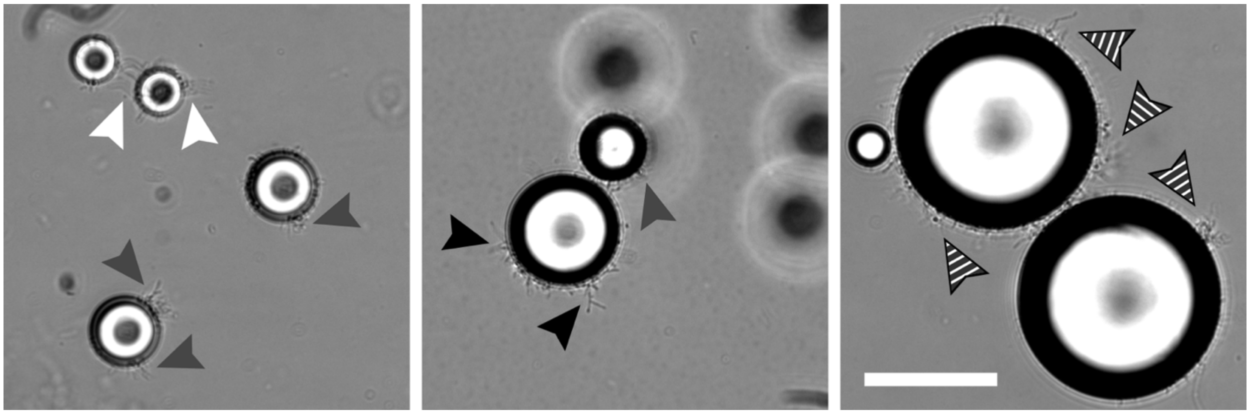

**Figure S7.** ***Sphingopyxis*** **sp. OPL5 cells attached to 15μm (white arrows), 25μm (gray arrows), 45μm (black arrows), and 75μm (striped arrows) diameter beads after 48 hours of incubation at 30˚C.** Scale bars = 50μm.

**Alt text**: Microscopy images of plastic beads and cells.

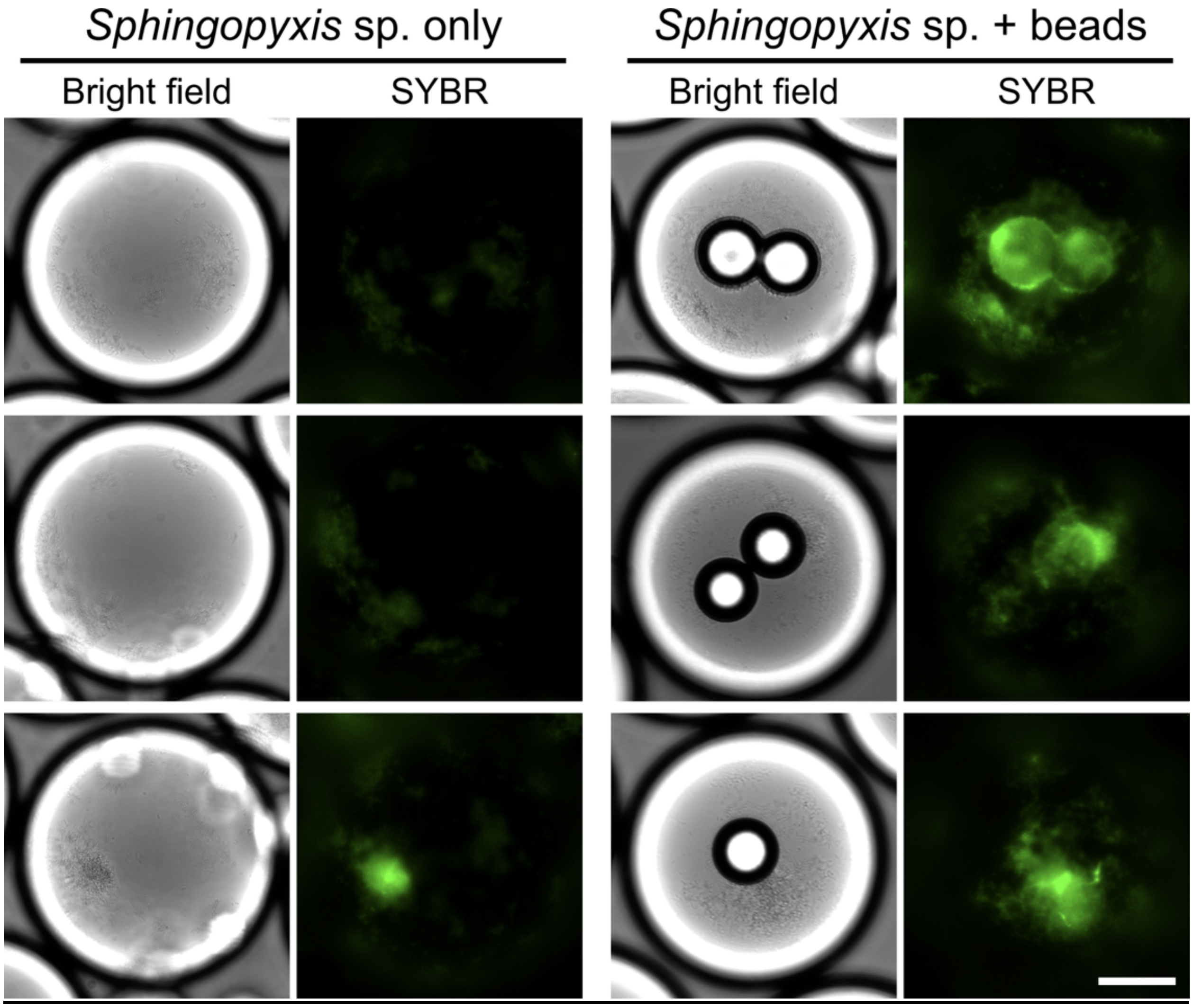


**Figure S8. Additional images of droplets containing *Sphingopyxis* sp. OPL5 with or without beads after 27-hour incubation.** Scale bar = 50μm.

**Alt text**: Zoomed-in microscopy images displaying cell attachment to and growth on the beads in droplets.

**Table S1**

PCR primers and expected fragment sizes.

| Target | Primer sequence (5' → 3') | Expected Length (bp) | Reference |
|---|---|---|---|
| 16S rRNA | F: AGAGTTTGATCCTGGCTCAG<br>R: GGTTACCTTGTTACGACTT | 1505 | (3) |
| *smf-1* | F: GGAAGGTATGTCCGAGTCCG<br>R: GCGGGTACGGCTACGATCAGTT | 82 | (2) |
| *rpfF* | F: CTGGTCGACATCGTGGTG<br>R: TGATCCGCATCATTTCATGC | 80 | (1) |
| *rmlA* | F: GCAAGGTCATCGACCTGG<br>R: TTGCCGTCGTAGAAGTACAGG | 151 | (1) |
| *spgM* | F: GCTTCATCGAGGGCTACTACC<br>R: ATGCACGATCTTGCCGC | 674 | (1) |
